# Human cytomegalovirus promotes *de novo* PC synthesis during early virus replication

**DOI:** 10.1101/2025.03.06.641752

**Authors:** Ian Kline, Rebekah L. Mokry, Yuecheng Xi, Magí Passols Manzano, Sidnie Layesa, Nowroz Sohrab Ali, Melissa A. Moy, Felicia D. Goodrum, John G. Purdy

## Abstract

Human cytomegalovirus (HCMV) infection reprograms metabolism, including lipid synthesis. While several metabolite-related pathways have been demonstrated to have altered activity in infected cells, the alteration of lipid-related pathways by HCMV has not been examined beyond fatty acid synthesis and elongation. In this study, we addressed this lack of understanding by focusing on phosphatidylcholine (PC), a class of lipids we previously showed is increased by HCMV infection in human foreskin fibroblasts. Here, we found that HCMV infection increases the abundance of PCs in several different fibroblasts and, similarly, in endothelial and epithelial cells. Additionally, HCMV elevates PC levels regardless of the level of confluency, type of growth medium, and presence of serum. Next, we investigated if HCMV alters the activity in the three PC synthesis pathways. We demonstrate that HCMV infection promotes the activity in the *de novo* PC synthesis pathway using a ^13^C-choline isotopic tracer and liquid chromatography high resolution tandem mass spectrometry (LC-MS/MS). Infection did not alter the activity in the other two pathways. Moreover, we examined the kinetics of PC remodeling by HCMV and found that the de novo synthesis pathway is promoted and the PC lipidome shifts 24 hours post infection. That led us to examining if the early stages of replication are sufficient to alter PC levels. After inhibiting late virus replication, we found that HCMV alters the PC lipidome independent of late gene expression. Overall, this work suggests that an immediate-early or early viral protein promotes the reprogramming of host lipid metabolism to ensure the synthesis of a lipidome necessary to support HCMV replication.

**IMPORTANCE:** Human cytomegalovirus (HCMV) is a common herpesvirus that establishes a lifelong and persistent infection in its human host (1). HCMV infection in most people does not cause overt disease (1). However, in immunocompromised individuals, severe CMV-associated disease can lead to permanent disabilities and even death (1, 2). Additionally, congenital CMV is the leading infectious cause of birth defects (3, 4).

Viruses have evolved to hijack host metabolic pathways to facilitate their replication cycle. In this study, we determine that HCMV promotes the activity in the de novo pathway of phosphatidylcholine (PC) synthesis. We demonstrate that the activity in the other PC synthesis pathways, the PEMT and Lands cycle, is unaltered by HCMV infection. Moreover, we found that HCMV infection alters metabolic activity to increase the PC lipidome before 48 hpi. Additionally, we demonstrate that changes in PC lipids during virus replication is independent of late gene expression. Together, our findings demonstrate that infection promotes the *de novo* PC pathway to increase PC lipids during the early stages of virus replication.

## Introduction

Human cytomegalovirus (HCMV) is a β-herpesvirus that causes disease in immunocompromised patients (1, 2) and developmental disabilities during congenital infection (3, 4). HCMV establishes life-long, persistent infection in part by reprogramming host metabolism to support virus replication. Infection promotes the activity in host lipogenic pathways for increased fatty acid (FA) synthesis and elongation to generate lipids needed for virus replication (5–8). Our previous lipidomic studies found that infection increases the relative abundance phosphatidylcholine (PC) (9, 10). PC is a class of phospholipids and a major constituent of biological membranes, including the virion envelope (11). PCs function in several biological contexts, including as structural and barrier lipids in lipid bilayers (11–14). The plasma membrane and many organelles are enriched with PC lipids, where they provide structural roles and help maintain basic cellular functions.

In humans, PC lipids are made by the *de novo* PC synthesis pathway (15–20). The de novo pathway is also known as the Kennedy pathway. In this pathway, choline and diacylglycerol are metabolized to generate PC. Two additional pathways can contribute to PC synthesis. The phosphatidylethanolamine N-methyltransferase (PEMT) pathway converts phosphatidylethanolamine (PE) lipids to PC lipids. This pathway is active in the liver where ~30% of PC is made via the PEMT pathway. Other tissues may have low levels of PEMT activity. The final pathway involves the addition of a free fatty acyl chain to a one-tailed lysophosphatidylcholine (LPC) to generate the two-tailed PC. Several lysophosphatidylcholine acyltransferases (LPCATs) can generate LPC to PC; this pathway is called the LPCAT pathway or Lands cycle. Given that HCMV infection increases the PC lipid content (5–11), we hypothesized that HCMV infection reprograms host metabolism to induce PC synthesis. We used metabolic tracing to measure the impact of HCMV infection on the activity of each of the three routes of PC generation. To this end, we used liquid chromatography high resolution tandem mass spectrometry (LC-MS/MS) to measure the lipid content and metabolic activity in primary human cells infected with HCMV. We determined that HCMV infection significantly altered the levels of several PC lipids, including PC with very long chain fatty acids (VLCFAs; PC-VLCFAs).

Here, we show that HCMV promotes PC and PC-VLCFA synthesis through metabolic reprogramming of the canonical *de novo* PC pathway at an early time post infection. Further, we found that increased PC abundance caused by infection are independent of late gene expression using an inhibitor of viral genome replication and LC-MS/MS. The findings suggest that an immediate early or early viral protein promotes host metabolic reprogramming that increases the *de novo* PC synthesis pathway and remodels the host lipidome.

## Results

### Lytic HCMV replication reprograms the PC lipidome

While several studies have demonstrated that lytic HCMV infection promotes lipid metabolism these studies have mostly investigated fibroblast cells grown under similar conditions (5–10, 21). We initiated our study by examining a diverse set of conditions to determine if several factors influence the ability of HCMV to alter PC lipid levels. We considered cell type, cell confluency, presence or absence of serum, and the cellular source of virus used to initiate infection. First, we used LC-MS/MS to measure the relative abundance of PC lipids in HCMV-infected cells and uninfected control human foreskin fibroblast cells (HFFs) that are fully confluent and serum-free, which have been extensively used to study metabolism in HCMV infection (5–7, 9, 10, 22). For these experiments, confluent cells were switched from containing fetal bovine serum (FBS) to serum-free one day prior to infection. We infected cells with TB40/E-GFP at a multiplicity of infection (MOI) of 3 in the absence of serum, and after a 1 h infection period, the cells were washed and returned to cell DMEM culture medium lacking FBS. We observed an overall increase in the relative abundance of most PC lipids 72 hours post infection (hpi), consistent with our previous observations (**Fig. 1A *column 1***) (9, 10).

**Figure 1:**
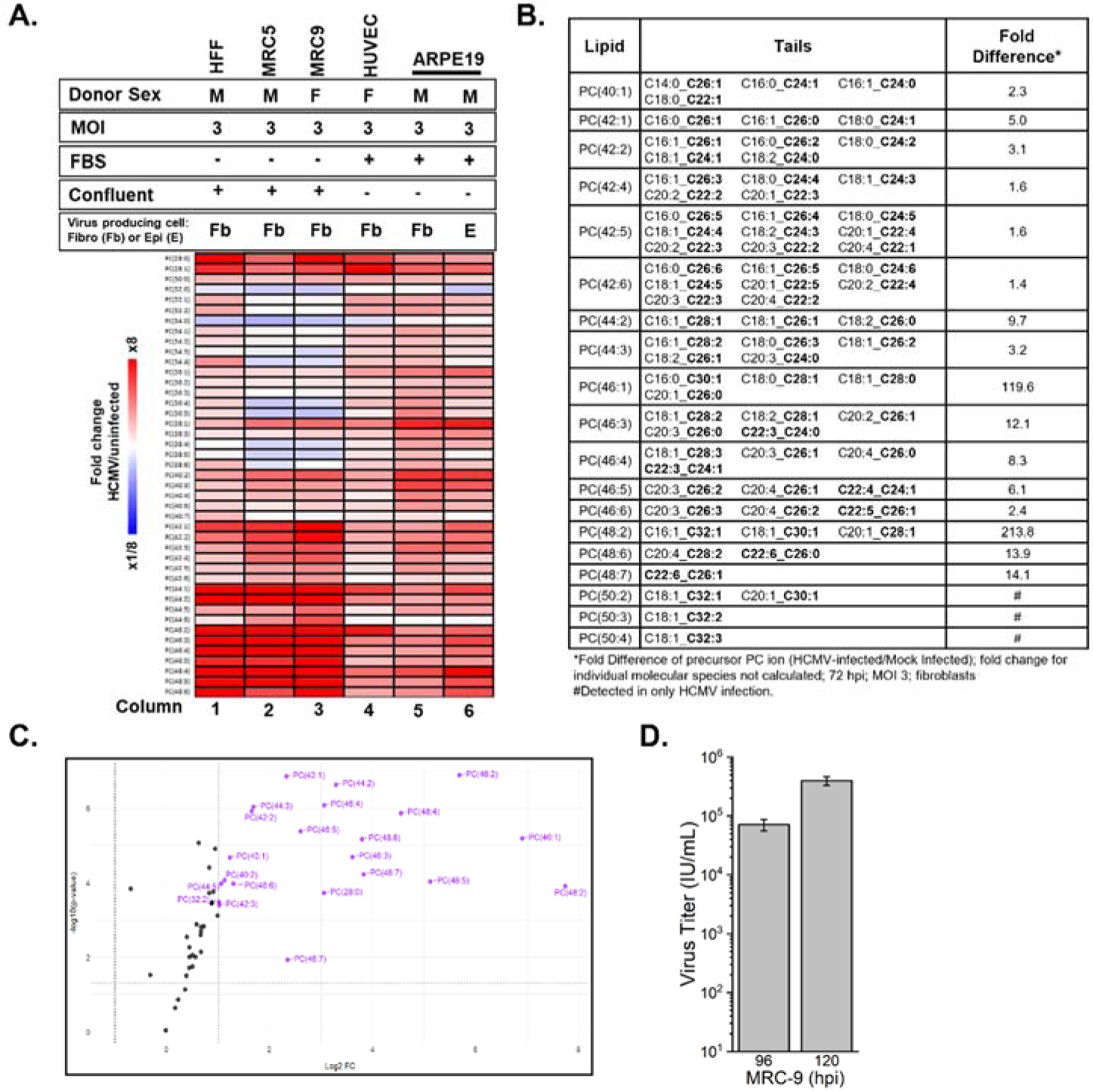
HCMV infection shifts the PC lipidome. **(A)** Heatmap of changes in the relative abundance of PC from HCMV-infected cells as measured by liquid chromatography high-resolution tandem mass spectrometry (LC-MS/MS). Cell types include fibroblast (HFF, MRC5, and MRC9), endothelial (HUVEC-TERT), and epithelial (ARPE-19) cells. Data were log transformed, ranked by increasing PC tail length, and represented as a relative abundance of infected to uninfected cells. In addition to cell type, different media, presence or absence of fetal bovine serum (FBS), confluence, type of cells used for generating virus stocks for infection, and multiplicity of infection (MOI) were tested. HCMV TB40/E-GFP was used for all infections. HFF N= 9, MRC5 N= 1, MRC9 N= 1, HUVEC N= 3, ARPE-19 N= 1. **(B)** Fold change in relative abundance of PC-VLCFA lipids. The fatty acyl tail composition is listed and VLCFAs are bolded. Data is from fully confluent, serum-starved HFF cells, N= 9. Significance was determined by t-test; P < 0.05. **(C)** Volcano plot showing changes in PC levels in HCMV-infected cells relative to uninfected cells. Fold change is log_2_ transformed and p-value is log10 transformed. PC lipids with ≥2-fold difference and P< 0.05 are purple with their name displayed. Significance was determined by t-test; P< 0.05. **(D)** HCMV replication in MRC-9 fibroblasts.

HCMV replication is supported by host fatty acid elongases that generate VLCFA lipids (7, 8). Previous studies demonstrated that HCMV infection increases the levels of fatty acids with 26 or more carbons (≥C26), with C30 being the longest identified tail in PCs (9, 10). Since we observed PC-VLCFAs increased by HCMV infection, we identified the VLCFA tail content of PC-VLCFA lipids in HCMV-infected cells. The longest VLCFA tails identified in HCMV-infected cells included C32:1, C32:2 and C32:3, demonstrating that HCMV infected cells have PCs with VLCFA longer than previously described. These tails were found in PC(50:2), PC(50:3), and PC(50:4), lipids that were readily detected in HCMV-infected HFFs and below the limit of quantification in uninfected cells (**Fig. 1B**).

We identified several previously unreported molecular species of PC-VLCFA containing either a saturated (no double bond) or monounsaturated fatty acid (MUFA; one double bond), paired with a long chain or very long chain polyunsaturated fatty acid (PUFA; two or more double bonds) such as C20:4 or C22:6 (**Fig. 1B**). PC(42:4), PC(42:5) and PC(42:6) differed from most others in their VLCFA tail content, each containing a previously unreported C26:3, C26:4, C26:5, or C26:6 tail that was paired with either a C16:0 or C16:1 tail (**Fig. 1B**). The levels of these PC-VLCFAs were ~1.5-fold higher than in uninfected cells (**Fig. 1B**). Overall, HCMV infection significantly altered the levels of PC lipids with ≥40 carbons (**Fig. 1C**). Together, these observations expand upon our understanding of the changes to PC lipid levels and their tail composition in HCMV-infected HFFs.

Next, we investigated if HCMV infection reprogrammed PC levels similarly in other fibroblast cell types that are grown in the same DMEM culture medium conditions. MRC-5 cells are another primary human fibroblast commonly used in HCMV metabolic studies (6, 8, 23, 24). Therefore, we measured the levels of PC from HCMV-infected MRC-5 and found increased PC-VLCFA lipid levels similar to HFFs (**Fig. 1A *column 2***). Next, we examined MRC-9 human fibroblast cells. While HFF and MRC-5 cells are from a male donor, MRC-9 cells were isolated from a female donor (25, 26). Like the other fibroblasts that have been well studied by the HCMV field, MRC-9 cells support HCMV replication (**Fig. 1D**). PC lipids in MRC-9 cells were remodeled by HCMV infection similar to the other fibroblast cells examined (**Fig. 1A *columns 1-3***)

HCMV exhibits broad cell tropism (27–30). Endothelial cells support HCMV infection *in vivo* (31–36). *In vitro* infection of endothelial cells supports lytic replication; however, the amount of infectious virus produced is lower than fibroblasts (34, 37, 38). We investigated the effects of HCMV on the level of PCs in subconfluent human umbilical vein endothelial cells (HUVECs) cultured in Endothelial Growth Medium (Lonza) with 10% FBS. At 96 hpi, HCMV-infected HUVECs had elevated levels of PCs (**Fig. 1 *column 4***). Again, PC-VLCFAs were increased by HCMV infection, albeit to a lesser degree than in fibroblasts.

Next, we examined PC lipids in epithelial cells which are infected *in vivo* and capable of supporting productive HCMV infection (27, 38). The retinal pigment epithelium (RPE) of the eye is a site of persistent HCMV infection leading to retinitis and blindness and retinal epithelial cells are a model for HCMV lytic replication (39–41) (42). We measured the levels of PC lipids in subconfluent ARPE-19 cells growing in 1:1 DMEM:F12 media containing 10% fetal bovine serum (FBS) infected with TB40/E at MOI 3 using a virus stock generated in fibroblast and titered on ARPE-19 cells to improve HCMV infection efficiency (42). At 72 hpi, HCMV-infected ARPE-19 cells contained a greater relative abundance of PC-VLCFA lipids than uninfected epithelial cells (**Fig. 1 *column 5***). In general, the level of most PC lipids was increased in ARPE-19 cells, including the PC-VLCFAs.

HCMV virus produced by epithelial cells can have a greater infection efficiency than virus produced by fibroblasts due altered ratios of glycoproteins present on the surface of the virion (43, 44). We tested if the type of cell producing virus alters the ability of HCMV to remodel PCs by infecting ARPE-19 cells with virus produced by fibroblast or ARPE-19 cells. A similar PC profile was observed in cells infected with virus produced in fibroblast and epithelial cells (**Fig.1 *columns 5-6***).

Collectively, these results demonstrate that HCMV lytic infection promotes the level of PC lipids independent of cell type, confluency, and culture medium. Further, these observations suggest that HCMV infection alters host metabolism in a broad range of cell types and growth mediums to increase the abundance of PC lipids.

### HCMV infection promotes *de novo* PC synthesis

Since HCMV infection promotes fatty acid synthesis and elongation (6–10) and the abundance of PC lipids (**Fig. 1**), we hypothesized that HCMV infection stimulates PC synthesis. PC *de novo* synthesis is the canonical pathway in humans (19). In this pathway, the headgroup comes from metabolism of choline while the tails are from diacylglycerol (DAG) (**Fig. 2A**). Choline is metabolized through a series of reactions to generate CDP-choline. In the final step, CDP-choline donates phosphocholine to (DAG, generating PC. To determine if HCMV infection alters metabolic activity in *de novo* PC synthesis, we measured PC synthesis in HCMV-infected and uninfected cells using ^13^C-choline stable isotope labeling. The labeled form of choline used contained two ^13^C atoms that will be retained at each step in the pathway resulting in PC lipids with two ^13^C atoms in the head (**Fig. 2A, red asterisk**). For these labeling experiments, HFFs were infected with TB40/E at a MOI of 3 for 1h. The experiments were performed in serum-free conditions, avoiding the potential for unlabeled choline from serum affecting the results. At 1 hpi, the virus inoculum was removed, the cells were washed twice, and DMEM containing ^13^C-choline in place of ^12^C-choline was added to the cells. From 24 to 72 hpi, lipids were extracted and the percent labeling in PC lipids was determined by LC-MS/MS following correction for the natural isotopic abundance. In uninfected cells, we observed an increase in the amount labeled PCs from 1.9-12.2% at 24 h to 13.7-31.4% at 72 h (**Fig. 2B**). In infected cells, we observed an increase in the percentage of ^13^C-labeled PC from 1.8-30.2% at 24 hpi to 33.3-53.6% at 72 hpi (**Fig. 2B**). At 24 hpi, none of the PCs showed a significant difference in labeling between infected and uninfected. Most of the identified PCs, 29 of 38, had a significantly greater percentage of ^13^C-labeling in HCMV-infected cells compared to uninfected cells at 48 hpi (**Fig. 2C**), demonstrating that HCMV infection promotes *de novo* PC synthesis between 24 and 48 hpi. By 72 hpi, 33 of the 38 measured PCs had a significantly higher percentage of labeling in infection, showing that HCMV promotes *de novo* PC synthesis from early to late times in virus replication (**Fig. 2B-C**).

**Figure 2:**
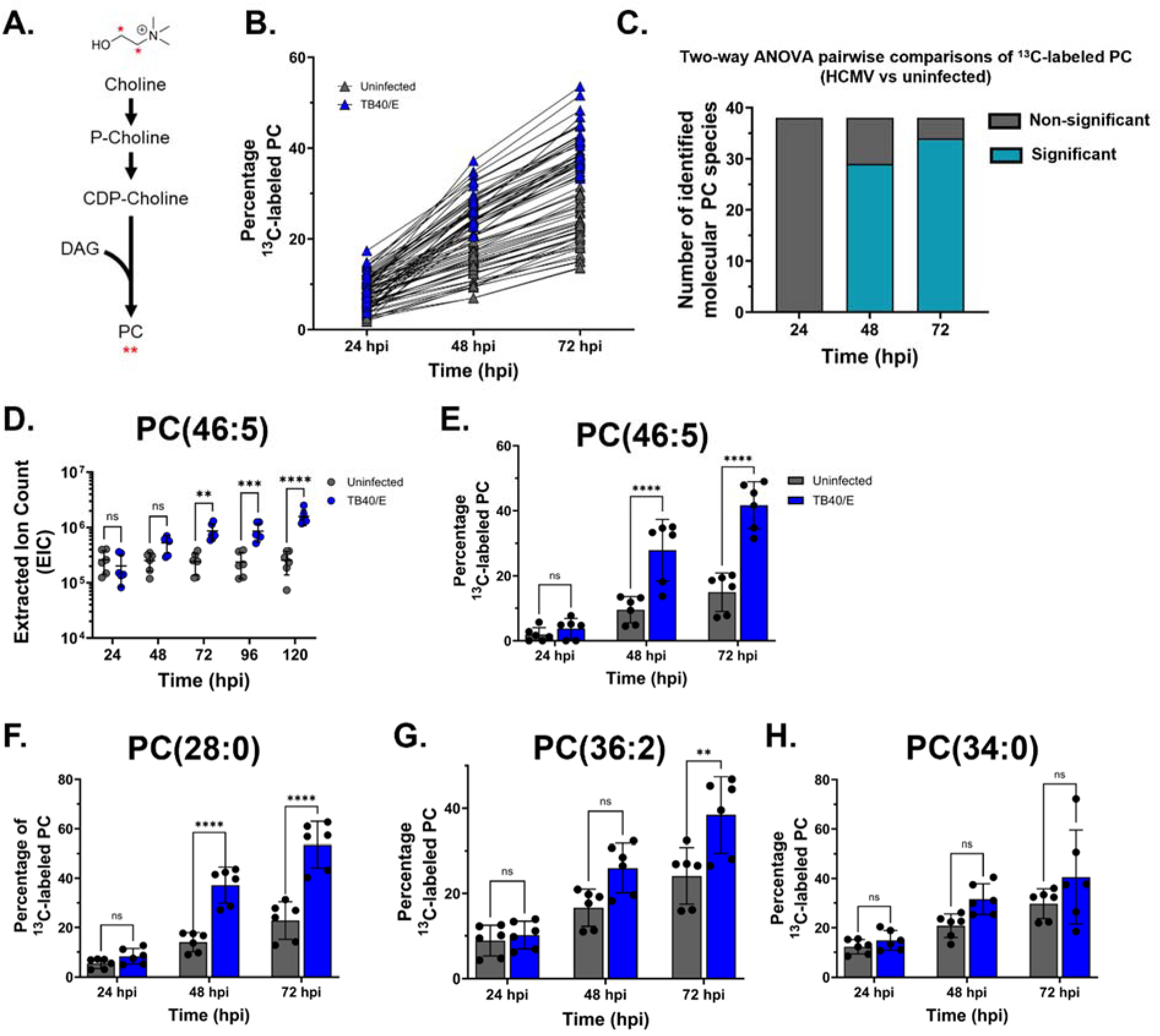
HCMV infection promotes metabolic activity in the de novo PC synthesis pathway. **(A)** The de novo PC synthesis pathway and ^13^C-choline metabolic labeling strategy. ^13^C carbons are denoted by red asterisks. HFF cells were grown to ful confluence, growth arrested for 3 d, and serum-starved for 24 hours. Cells were infected at an MOI 3 with HCMV strain TB40/E-GFP. Cells were cultured in medium containing ^13^C-choline starting at 1hpi. **(B)** *De novo* PC pathway activity in HCMV-infected and uninfected cells. **(C)** Number of significant pairwise comparisons between HCMV-infected and uninfected cells (38 lipids identified in total). **(D)** Ion count values from extracted ion chromatogram (EIC) showing the relative abundance of PC(46:5). **(E)** Percentage of 13C-labeled PC-VLCFA, PC(46:5). **(F-H)** Representative lipids PC(34:0) **(F)**, PC(36:2) **(G)**, and PC(28:0) **(H)** demonstrating observed pairwise phenotypes; (ns/ns/ns = 4), (ns/ns/sig. = 5) and (ns/sig./sig/ = 29). **(C-H)** Two-way ANOVA, with Tukey’s test, P < 0.01 **; P < 0.001, ***; P < 0.0001, ****. N= 3

Next, we selected PC(46:5) as a representative PC-VLCFA to highlight the changes over time that we observed in the *de novo* synthesis pathway. By 72 hpi, its abundance was increased and the level remained elevated through 120 hpi in HCMV infection (**Fig. 2D**). We mimicked infection by feeding uninfected cells ^13^C-choline for 71 h and found that 15% of PC(46:5) was labeled (**Fig. 2E**). In HCMV-infected cells, more than 40% of PC(46:5) were labeled. These data demonstrate HCMV infection increases the relative abundance of PC(46:5) and promotes its synthesis via the *de novo* pathway suggesting that HCMV promotes the synthesis PC-VLCFAs to increase their levels in infection.

Since we observed an increase in the labeling of most PC lipids, PC-VLCFAs and those with shorter tails (**Fig. 2B**), our results suggest that HCMV infection promotes the *de novo* synthesis of PCs regardless of tail composition. To investigate this possibility further, we examined lipids with labeled percentages that were significantly increased at 48 and 72 hpi, similar to PC(46:5). Several of these lipids contained VLCFA tails or shorter tails, including PC(28:0), the shortest PC measured in our study (**Fig. 2F**). Of the 38 lipids we identified, 29 had the same phenotype, including PC lipids ranging from PC(28:0) to PC(46:5) demonstrating that HCMV infection increases the *de novo* synthesis of PCs regardless of tail type. For five PCs, labeling in infected cells was increased only at 72 hpi, for example PC(36:2) (**Fig. 2G**). The others were: PC(32:0), PC(40:8), PC(42:4), and PC(42:6). Only 4 PCs had similar percentage labeling in uninfected and HCMV-infected cells from 24-72 hpi: PC(32:2), PC(34:0), PC(36:3), and PC(46:2)(**Fig. 2H**). Overall, our ^13^C-choline labeling results demonstrate that HCMV infection promotes the activity of the *de novo* pathway for most PC lipids regardless of tail composition, suggesting that this pathway contributes to an increase in the relative abundance of many PC lipids.

### The PEMT pathway is inactive in fibroblasts and unaffected by HCMV

In addition to the *de novo* synthesis pathway, PC lipids can be synthesized via the phosphatidylethanolamine *N-*methyltransferase (PEMT) pathway. PEMT is a host enzyme that adds three methyl groups to the head of PE to generate PC (**Fig. 3A**) (19, 45–49). PEMT expression is greatest in the liver, epididymis, and adipose tissue with hepatocytes and a few other cell types using PEMT to generate part of their PC content (50, 51). We investigated if HCMV infection alters the expression of PEMT or activity in the pathway. First, we examined if PEMT protein is present in HFF cells. PEMT protein levels in uninfected primary human fibroblasts were undetectable (**Fig. 3B**), consistent with the known expression of PEMT (45, 52). Next, we examined if HCMV infection alters PEMT levels in fibroblasts. Similar to uninfected cells, PEMT protein was unobservable in HCMV-infected fibroblasts up to 48 hpi (**Fig. 3B**). When blotting for PEMT, we observed a 25 kDa band in infected samples at 72 and 96 hpi, however a band at the expected 19 kD size was not detected (**Fig. 3B**). We evaluated the PEMT antibody by exogenously expressing PEMT using a doxycycline inducible system and used the same system to express GFP as a control. When we expressed PEMT in HFF cells, we observed a band at the expected size of 19 kD ensuring that our western blot approach was able to detect PEMT and that it migrates at the anticipated size (**Fig. 3B**). PEMT has two isoforms, PEMT-S and PEMT-L (53). PEMT-S has higher enzymatic activity than PEMT-L. PEMT-S is approximately 19 kDa and PEMT-L can be N-glycosylated and appears as a band at 24-28 kDa (53). We treated HCMV-infected samples with PNGase F to determine if the 25 kDa observed at late times in HCMV-infection represents glycosylated PEMT. PNGase F treatment had no effect on the 25 kDa band observed late in infection using the PEMT antibody (**Fig. 3C**). As expected, PNGase F treatment reduced the size of HCMV gB which is known to be glycosylated. The 25 kDa band may be unglycosylated PEMT-L or due to cross-reactivity with a late HCMV protein or a host protein that is only expressed late in HCMV replication. We conclude that HCMV infection does not lead to the expression of PEMT-S but may possibly cause the expression of PEMT-L during the late steps of replication.

**Figure 3:**
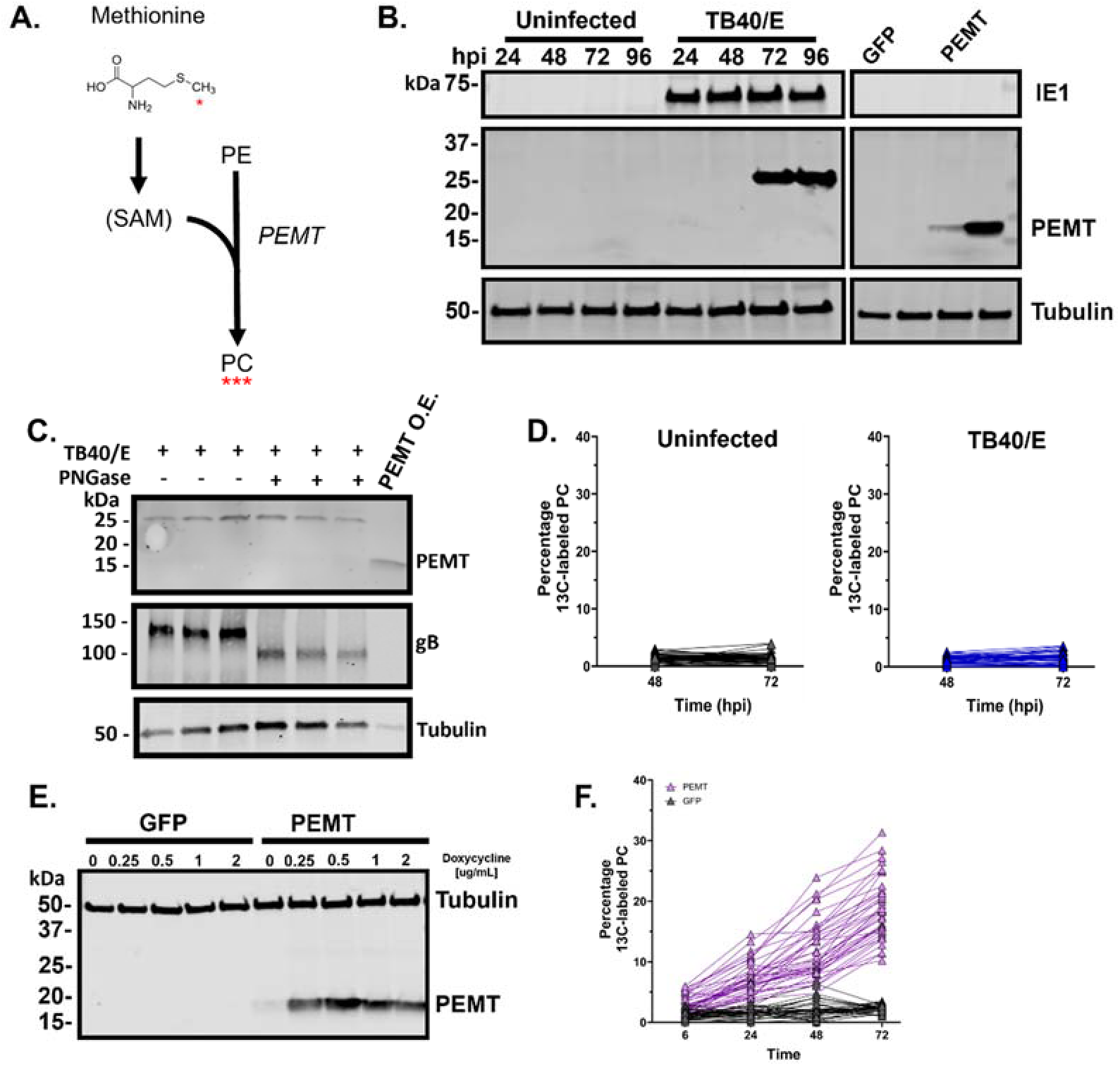
PEMT PC synthesis is not active in fibroblast cells. **(A)** The PEMT pathway and labeling strategy. Methionine is converted to S-adenosylmethionine (SAM). SAM is a methyl donor used for the sequential methylation of PE resulting in PC synthesis. ^13^C-methionine was used to measure PEMT; red asterisk denotes the labeled carbon. **(B)** Western blot for PEMT protein from infected and uninfected cells. HFF cells engineered to overexpress GFP and PEMT. Overexpression was induced using 1µg/mL of doxycycline. **(C)** Immunoblot of uninfected and TB40/E-infected HFF cells treated with PNGase F. Uninfected cells overexpressing PEMT or GFP were included. **(D)** ^13^C-methionine labeling of PC lipids. HFF cells were grown fully confluent, growth arrested, and serum-starved for 24 hours. Cells were infected at an MOI of 3 with HCMV strain TB40/E. At 1 hpi, cells were washed to remove unlabeled methionine and fed medium containing 13C-methionine. **(E)** Immunoblot of HFF cells expressing either GFP or human PEMT under control of a doxycycline inducible promoter (1 µg/mL doxycycline). **(F)** ^13^C-methionine labeling from cells overexpressing PEMT or GFP treated with 1 µg/mL doxycycline. 6, 24, 48 hpi n= 1, 72 hpi N= 2.

Since PEMT-L may potentially be expressed after 48 hpi, we sought to directly test if HCMV infection alters the activity in the PEMT pathway. The methyl donor used in the PEMT reaction is S-adenosylmethionine (SAM). SAM is generated from methionine (**Fig. 3A**). To measure PEMT activity, we fed cells ^13^C-labeled methionine and measured labeled PC lipids using LC-MS/MS. At 1 hpi, cells were washed prior to the addition of culture medium that contained ^13^C-methionine. We first measured the amount of ^13^C-lableled PC lipids in uninfected cells. We found the percentage of ^13^C-labeled PC ranged between 0.07- to 4% for any given species, indicating primary HFFs exhibit little to no PEMT activity (**Fig. 3D**). This is consistent with the lack of observable PEMT-S and PEMT-L enzyme in these cells. When we tested if HCMV infection altered PEMT activity, we observed a low level of ^13^C-labeled PC similar to that of uninfected cells (**Fig. 3D**). Our results indicate that PEMT activity is unaltered by HCMV infection.

Next, we ensured that our labeling strategy could measure activity in the PEMT pathway if it was active in HFF cells by generating PEMT-expressing HFF cells using a doxycycline inducible system (**Fig. 3E**). Feeding ^13^C-methionine to these cells resulted in 11- to 35% labeling in PCs when PEMT expression was induced with 1µg/mL of doxycycline (**Fig. 4F**). In contrast, control cells with doxycycline-inducible GFP expression had PC labeling percentages of 0.32- to 3.5%, levels similar to uninfected and HCMV infected fibroblasts (**Fig. 3F**). Based on the data presented here, we conclude that the increase in the abundance PC at 72 hpi following HCMV infection occurs independent of the PEMT synthesis pathway.

**Figure 4:**
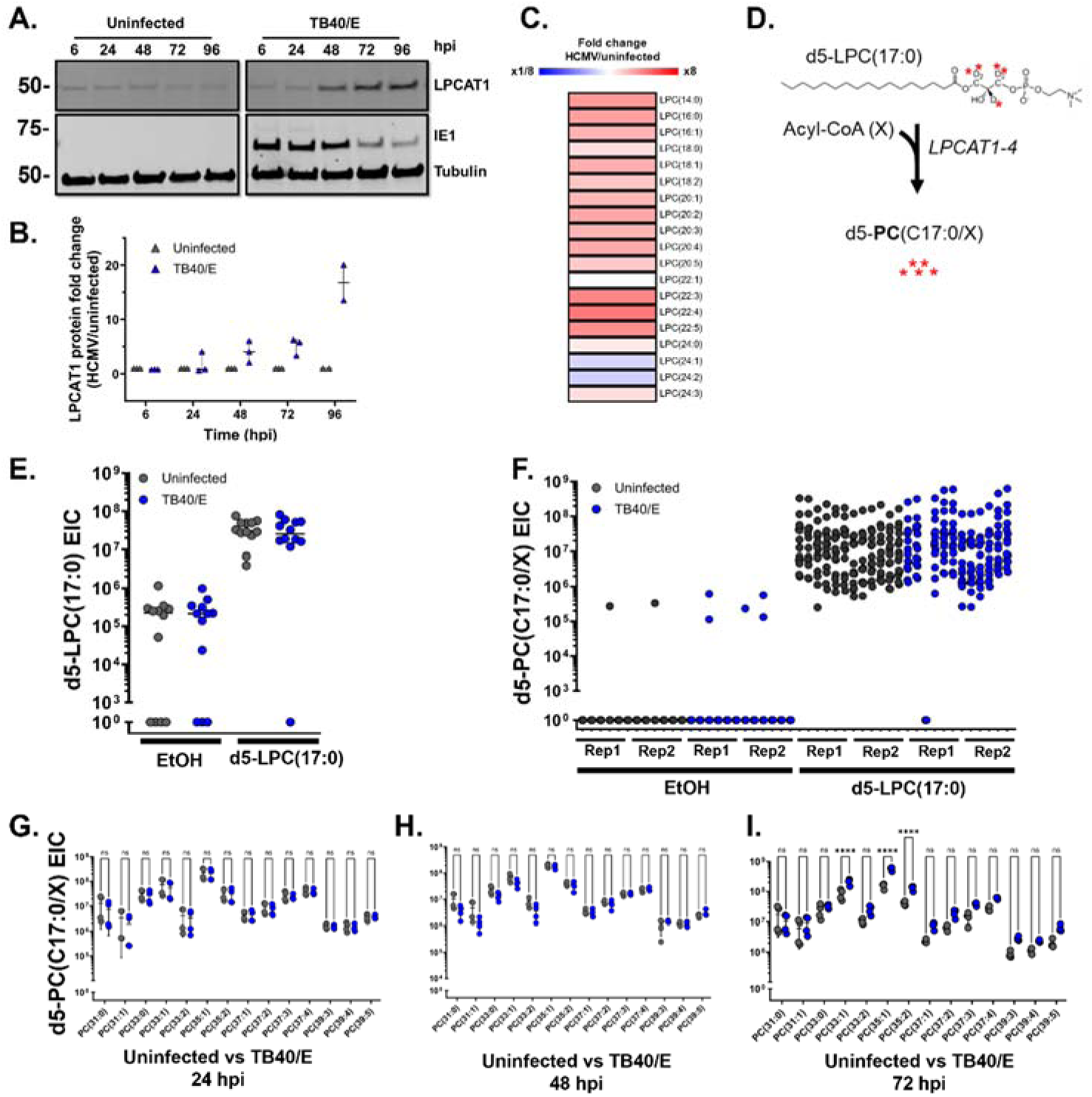
LPCAT is constitutively active in HCMV infection. **(A and B)** LPCAT protein levels were measured **(A)** and quantified **(B)** in uninfected and HCMV-infected cells. Representative blot from three biological replicates. HFF-hTERT cells were grown to full confluence, growth arrested for 3 d, and serum-starved for 24 hours. Cells were infected at an MOI of 3 with HCMV strain TB40/E-GFP. **(C)** LPC lipid abundance in HCMV-infected cells relative to uninfected cells, 72 hpi. **(D)** LPCAT pathway converts a one-tailed lysoPC (LPC) to a two-tailed PC by the addition of a free fatty acyl chain. To measure the activity in the pathway an exogenous deuterated 5-labeled LPC with a C17:0 tail [d5-LPC(17:0)] was used to monitor its conversion to a labeled PC that contains an additional tail at the sn2 position [d5-PC(17:0/X], where X is the newly added tail. Deuterium atoms in the glycerol backbone are denoted by red asterisks. **(E)** HFF-hTERT cells were grown fully confluent and serum-starved for 24 h prior to infection with TB40/E at MOI 3. At 1 hpi, cells were washed twice with PBS and cultured in medium containing either 10µM d5-LPC(17:0) or EtOH vehicle control. Levels of d5-label from d5-LPC(17:0) and EtOH vehicle treated cells. **(F)** Conversion to d5-PC(17:0/X) from HCMV-infected and uninfected cells treated with either 10 µM d5-LPC(17:0) or EtOH vehicle control. (G-I) d5-PC(17:0/X) lipids from HCMV-infected and uninfected control cells, 24 **(G)**, 48 **(H)** and 72 hpi **(I)**. P= 0.0001, ***; P< 0.0001 ****. N=2.

### LPCAT activity is unaltered by HCMV infection

A third route of synthesizing PC involves the generation of two-tailed PCs from one-tailed lysophosphatidylcholine (PC). In this pathway, one tail of the PC comes from LPC and the other from a free fatty acyl chain (54). The conversion of LPC to PC contributes to the maintenance of biological membranes by regulating PC composition through the activity of a family of host enzymes called lysophosphatidylcholine acyltransferase (LPCAT1-4) (54–58). Of these, LPCAT1 is reported to generate saturated PCs like those increased by HCMV infection (**Fig. 1A**) (9, 10).

First, we measured protein levels of LPCAT1 to initiate an investigation of the impact of HCMV-infection on the LPCAT pathway. A low level of LPCAT1 was observed in uninfected samples that remained at a similar level throughout the 6-96 hpi time course examined (**Fig. 4A**). Beginning at 48 hpi, LPCAT1 started to rise in HCMV-infected cells (**Fig 4A and B**). At later times, the LPCAT1 was elevated in HCMV infection. Next, we measured the levels of LPC at 72 hpi. We found that LPC levels were generally increased by HCMV infection (**Fig. 5C**).

**Figure 5:**
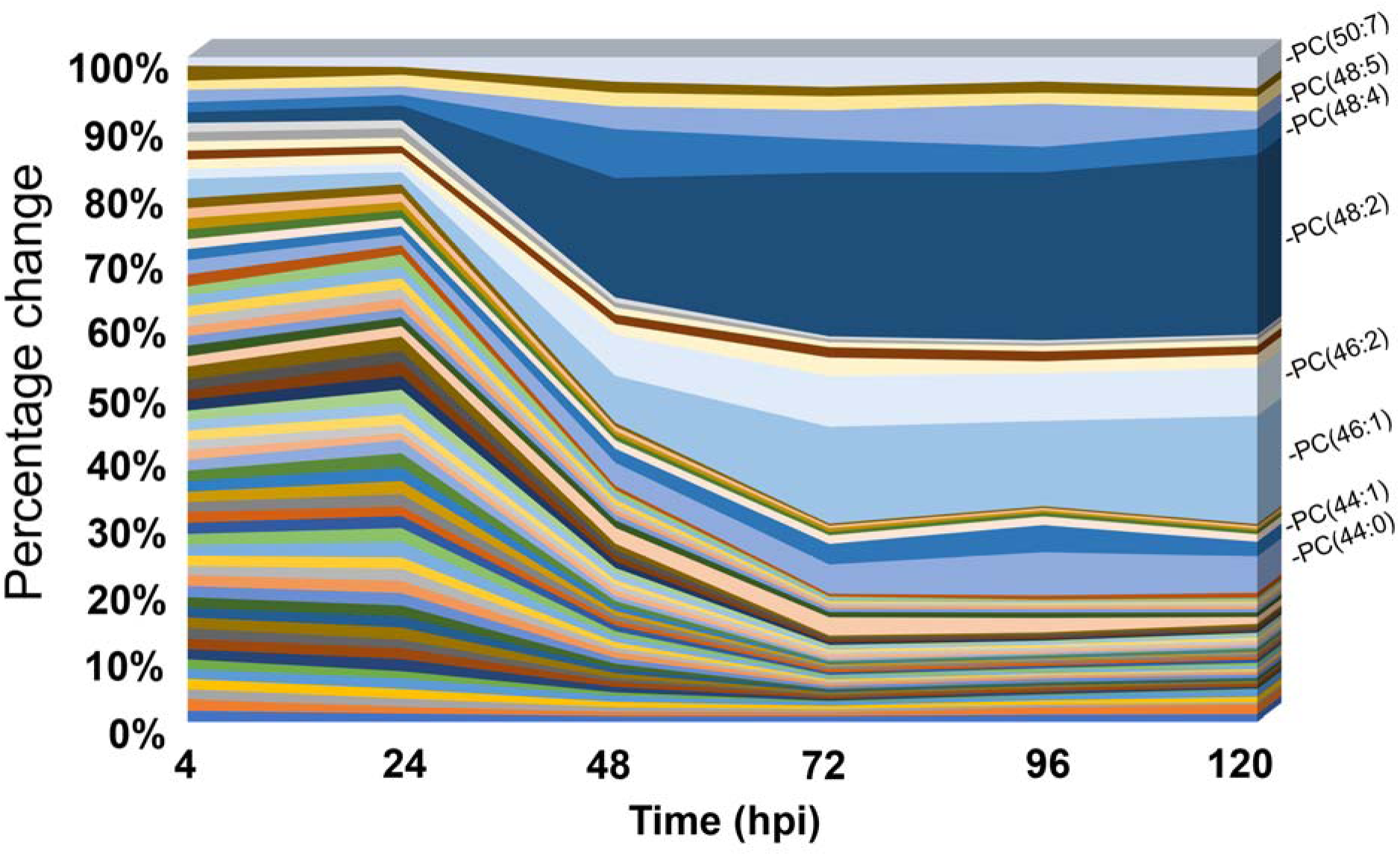
HCMV alters the PC lipidome starting at 24 hpi. The fold-change (infected relative to uninfected) in each PC was determined and expressed as a percentage of the sum total of the fold-change for all PC lipids. Confluent HFF cells were serum-starved, infected with TB40/E-GFP at MOI 3. Lipids were extracted from 4-120 hpi and measured by LC-MS/MS. The PC lipids most altered by HCMV infection 120 hpi included; PC(44:0), PC(44:1) PC(46:1), PC(46:2), PC(48:2), PC(48:4), PC(48:5), and PC(50:7). N=3.

To test if HCMV-infection impacts LPCAT activity, we fed cells a deuterium (d) labeled LPC lipid and measured labeled PCs using LC-MS/MS (**Fig. 5D**). Deuterium-labeled d5-LPC(17:0) contains a single odd-chain saturated C17:0 FA and five deuterium atoms located in the glycerol backbone that generate a combined mass shift of Δ5.0313 *m/z* higher than endogenous unlabeled LPC(17:0). In this assay, we used LC-MS/MS to track the conversion of d5-LPC(17:0) to d5-PC(17:0/X), where X represents the addition of a complementary fatty acyl-CoA that was added to the *sn2* position by an LPCAT (**Fig. 5D**). First, we determined the background levels of d5-LPC(17:0) and uptake of the fed exogenous d5-LPC(17:0) substrate in HCMV-infected and uninfected cells. The d5-LPC(17:0) substrate was in a solution of ethanol (EtOH) and was fed to cells using 1.7 µM BSA in a final concentration of 0.5% EtOH. In cells treated with 10 µM d5-LPC(17:0) we detected a signaling that was >100-fold greater than background signal demonstrating that uninfected and infected cells take up the fed LPC at a similar level (**Fig. 4E**). To evaluate LPCAT activity, we compared the conversion of d5-LPC(17:0) to d5-PC(17:0/X). First, we examined the background levels of d5-PC(17:0/X) in EtOH only treated cells. We observed that only 7 of 336 peaks in EtOH treated samples, demonstrating that uninfected and HCMV-infected cells have little to no endogenous levels of d5-PC(17:0/X) (**Fig. 4F**). Since natural d5-PC(17:0/X) levels are low, we report the extracted ion counts instead of percentage of labeling In d5-LPC(17:0) treated cells, d5-PC(17:0/X) lipids were readily identified demonstrating that LPCAT conversion of d5-LPC(17:0) to d5-PC(17:0/X) is active in these cells (**Fig. 4F**). Moreover, d5-PC(17:0/X) levels were similar in uninfected and infected cells suggesting that LPCAT activity is unaffected by HCMV infection (**Fig. 4F**).

Since we found that the *de novo* PC synthesis pathway was increased by HCMV infection starting at 24 hpi (**Fig. 2B**), we measured LPCAT activity via conversion of d5-LPC(17:0) to d5-PC(17:0/X) at 24-72 hpi. At 24 and 48 hpi, a similar level of labeled PC in uninfected and HCMV-infected cells was observed in all identified d5-PC(17:0/X) lipids(**Fig. 4G and H**). At 72 hpi, we found that only 3 of the 14 d5-PC(17:0/X) lipids had a statistically higher amount of labeled PC in infected cells (**Fig. 4I**). For these three, the difference between infected and uninfected was between 2.2 and 3.9-fold increased. Overall, these findings show that the LPCAT pathway is active in primary human fibroblasts and that HCMV infection has little to no impact on the pathway activity.

### HCMV reprograms host PC lipid metabolism during the early stage of replication

Thus far, we have found that HCMV infection increases the abundance of PCs and their *de novo* synthesis starting at 24 hpi. Next, we wanted to define the time when infection alters PC levels. Therefore, we measured the relative abundance of PCs in HCMV-infected and uninfected cells from 4-120 hpi. In this case, we determined the fold-change for each individual PC and expressed the percentage change where the total fold-change adds up to 100. In this case, if 100 PCs were measured and all had the same change then each would have 1% change. At 4 hpi, we observed a similar change in abundance among all PC lipids, each individual PC accounted for <2% (**Fig. 5**). At 24 hpi, the percentage change of most PCs were similar to 4 hpi. Between 24 and 48 hpi, the levels of some PC-VLCFA lipids, including PC(44:0), PC(44:1), PC(46:1), PC(46:2), PC(48:2), PC(48:4), PC(48:5), and PC(50:7), rose precipitously, accounting for the majority of change observed (**Fig. 5**). At the later times, 72-120 hpi, these PC-VLCFAs were the PCs that changed the most in HCMV infection. These data support that the PC lipidome is remodeled during the early stage of HCMV replication starting after 24 hpi and continues to be shifted throughout the later steps of replication.

Based on our findings that PC lipids levels and their *de novo* synthesis are increased starting around 24 hpi, we hypothesize that the PC lipidome is altered independent of late genes. HCMV genes are classified broadly into three major kinetic classes, immediate early (IE), early (E), and late (L). Since late gene expression requires viral genome synthesis (59), we used phosphonoacetic acid (PAA) to inhibit HCMV genome synthesis and block the expression and synthesis of late genes (60). Fibroblast cells were infected with TB40/E and treated with PAA or vehicle (H_2_O) at 1 hpi. PAA was readministered every 24 h. PAA treatment reduced virus genome synthesis (**Fig. 6A)**. PAA treatment reduced the levels of late proteins pp28 and pp71, while IE1 protein and early protein pUL44 were not impacted during the first part of replication (**Fig. 6B-F**). A reduction in immediate-early and early proteins were observed at later times, as previous shown (59, 61–63). Since PAA treatment was inhibiting genome synthesis and late protein expression as expected, we investigated the impact of PAA treatment on PC lipid levels. At 72 and 96 hpi, the relative abundance of PC lipids was measured by LC-MS/MS. PAA treatment had little effect on PC levels (**Fig. 6G**). These findings demonstrate that HCMV infection alters the PC lipidome independent of viral genome synthesis and late protein expression.

**Fig 6:**
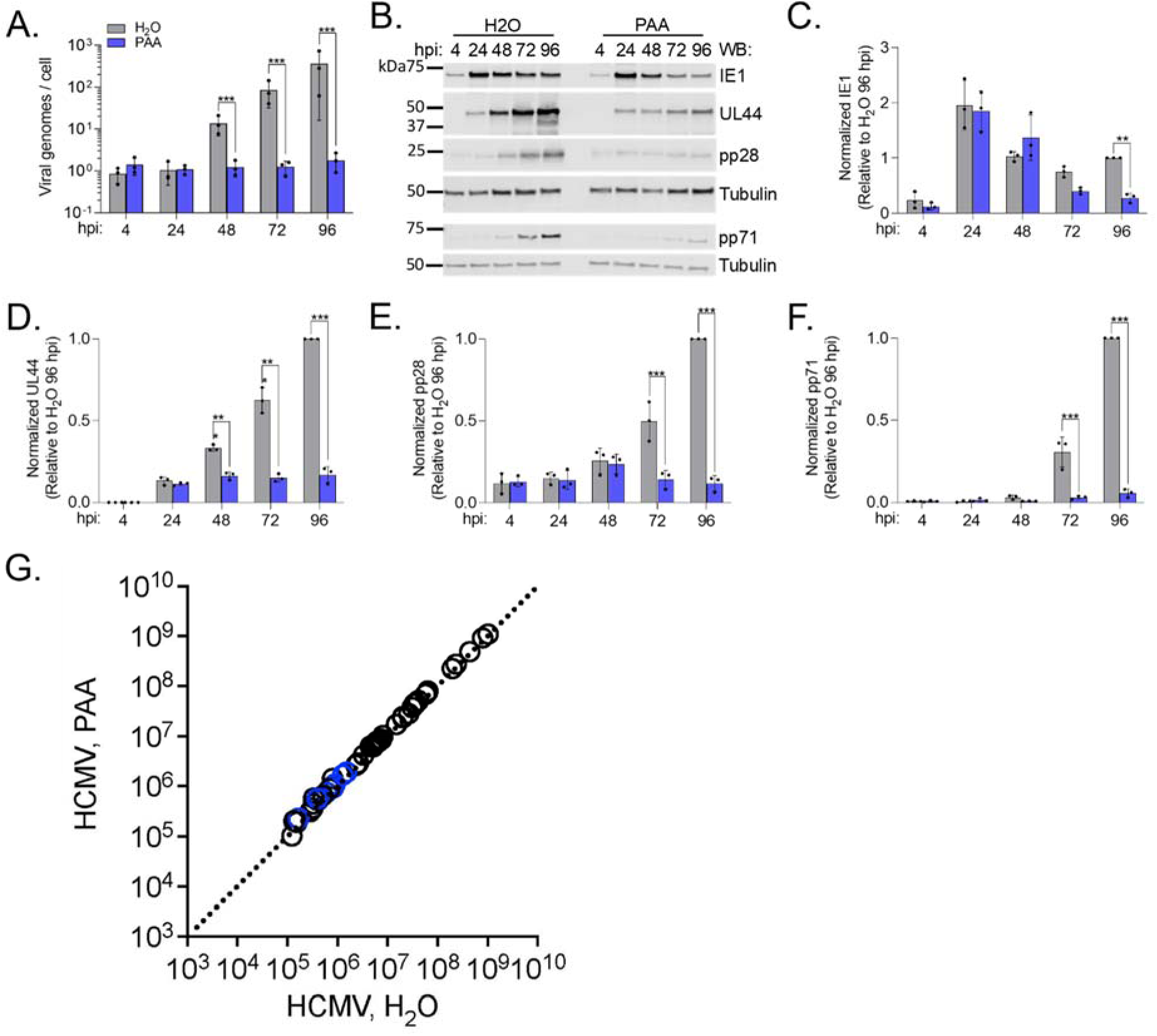
Increased PC synthesis is not dependent on viral genome synthesis. HFFs were grown to confluence, growth arrested for three days, and serum starved for 24 hrs. Cells were infected at MOI 3 with TB40/E-GFP. At 1 hpi, cells were treated with 100 ug/mL phosphonoacetic acid (PAA) or H2O vehicle control. Treatment was replaced every 24 hpi. **(A)** Viral genomes per cells were determined using qPCR. **(B)** Whole cell lysates were collected and analyzed by western blot. Representative blot from three biological replicates. **(C-F)** Viral protein levels were normalized to tubulin and quantified relative to 96 hpi, H2O control. **(G)** Relative PC levels in HCMV-infected cells treated with PAA or H2O vehicle control. PCs were measured by LC-MS/MS following normalization to cell number. PC highlighted in Figure 1B are colored blue. **(A-F)** Statistics were performed on transformed data for A. Two-way ANOVA with Šídák’s test were used to determine significance. P < 0.01, **; P < 0.001, ***. N=3

## DISCUSSION

Multiple studies demonstrated that HCMV replication depend on increased lipid metabolism (6, 8–10, 22, 64). It is also known that the changes in metabolism occur temporally (5, 9, 65, 66). Since virus replication depends on changes in host metabolism, the temporal changes in metabolism may need to align with the kinetics of virus replication to generate an environment to support successful replication. Our current understanding of the temporal control metabolism by viruses is limited. Identifying the temporal control for virus-induced changes to lipid metabolism will guide studies in defining the viral proteins, including their kinetic classes, involved in metabolic reprogramming. In this study, we utilized a lipidomics approach to identify the synthesis pathway HCMV targets to increase PC lipids and defined when the activity in the pathway is altered.

Our work demonstrated that lytic infection of HCMV increases the levels of PC lipids independent of cell type, presence or absence of serum, and the cell type which the inoculating virus was produced (Fig. 1). Similarly, the metabolome is altered by lytic infection of HCMV and herpes simplex virus type 1 (HSV-1) independent of cell type (66). Moreover, PC-VLCFAs were among the lipids increased most in HCMV infection (Fig. 1) (9, 10). HCMV-induced FA elongation results in VLCFA tails that are present in membranes (8). While each type of cellular membrane has a distinct lipid composition, PC lipids are a major constituent of most membranes and are the most abundant phospholipid in several cellular membranes. Nearly half of the phospholipids in the double-layered membrane of the nucleus are PC (67), while the plasma membrane PC is 20-25% of the total lipid content and enriched on the outer monolayer (68). Viruses remodel host membranes to support their replication. For example, poxviruses protect the site of viral DNA replication by rearranging ER-derived membranes to form a mini-nuclei in the cytoplasm called virosomes or virus factories (69). HCMV genome replication occurs in the nucleus and assembling virus particles acquire their envelope in the cytoplasm during secondary envelopment. Their envelop membrane consists of 34% PC, 48% PE, 8% PS, 3% PI, and the remainder is composed of select PA and PG species (11). The lipid composition of the virion differs from the nuclear and plasma membrane and more closely resembles synaptic vesicles, suggesting viruses create a unique lipid environment that is used during secondary envelopment (11). Following secondary envelopment, virions must traffic out of the cells in a process that likely involves different lipids. Recent research demonstrated that HCMV egress is cell type dependent (70, 71). In addition to lipids supporting assembly, egress, and the virus envelop, the characteristic cytomegaly, and expansion of plasma and nuclear membranes observed in HCMV-infected cells suggest a necessary increase in the levels of lipids.

In this study, we found that PC metabolism was altered by HCMV independent of the expression of the late temporal class of genes. Herpesvirus lytic replication is coordinated by a well-regulated cascade of gene expression. Conventionally, viral gene expression during lytic replication is characterized as three kinetic classes: immediate-early, early, and late. Immediate-early genes are transcribed independent of any de novo viral protein translation (72). These genes include immediate-early genes 1 and 2 (IE1 and IE2) encoded by UL123 and UL122 genes. IE1 and IE2 proteins are transcriptional activators that coordinate the expression of viral genes. Early (E) genes require protein translation to occur after infection but do not require viral genome replication. HCMV encodes more than 100 E genes. The expression of late (L) genes depends on viral genome synthesis. Herpesvirus genes can be further classified as genes with transcription enhanced by viral DNA synthesis (delayed-early or leaky-late) and genes with transcription enabled by viral DNA synthesis (true-late). A recent RNA-seq study led to the proposal of classifying HCMV genes into seven temporal classes (TC1-7) (62). In this system, IE genes are classified at TC1, genes with early or delayed early kinetics as TC2-3, and late genes in TC5-7. TC4 contained genes that were expressed with early kinetics but had a dependency on DNA replication. The identification of leaky-late, true-late, and TC4-7 genes involves the use viral DNA replication inhibitors such as PAA or phosphonoformate (PFA). In this study, we treated HCMV infected cells with PAA to reduce viral genome replication and the levels of late proteins (Fig. 6A-F). We used this as a tool to determine that HCMV infection increased the levels of PC independent of viral DNA replication and late gene expression (Fig. 6G). These findings suggest that IE and E proteins are sufficient to promote changes in metabolism that result in PC increases during HCMV infection. While our findings demonstrate that late gene expression is unnecessary for PC changes that occur in infection, late gene expression may help promote metabolic changes started earlier in virus replication or may support other changes in the host lipidome.

Several viral proteins are involved in HCMV metabolic reprogramming. Of these, UL38 is the one we understand the most. Consistent with our conclusion that PC metabolism is altered with early kinetics, UL38 is an early gene. It is expressed within the first 24 h of infection in fibroblasts (73). UL38 suppresses cell death while triggering and maintaining mTOR activity (73, 74). UL38 supports fatty acid elongation that is necessary for virus replication, and it was originally proposed that HCMV does so through both mTOR-dependent and independent mechanisms (8). Subsequently, it was demonstrated that UL38 supports HCMV replication by inducing several additional metabolic activities, including increasing the consumption of glucose and some amino acids (75). While UL38 targeting of TSC2 also drives UL38 induction of mTOR activity, UL38 modulates metabolism independent of mTOR (75). Furthermore, UL38 inhibition of TSC2 aids its metabolic control. In addition to UL38, UL37x1 supports HCMV remodeling of the host lipidome, suggesting that it facilitates metabolic reprogramming (9). UL37x1 helps promote the expression of proteins involved in fatty acid elongation that use glycolytic-derived metabolites increased by UL38, suggesting that UL38 and UL37x1 may function in a coordinated fashion to alter metabolism. UL13 is the third HCMV protein found to be involved in metabolism (76). UL13 localizes to the mitochondria where it promotes oxidative phosphorylation. Since UL38, UL37x1, and UL13 are expressed early in lytic replication in fibroblasts (62), they can allow for viral-driven metabolic changes to occur prior to viral genome replication. The observation that the known viral proteins exerting metabolic control—UL38, UL37x1, and UL13—are expressed from early genes further supports our conclusion that HCMV reprograms metabolism in the early phase of replication to increase PC lipids.

Additional evidence in the host supports the conclusion that metabolic reprogramming is initiated during the initial stages of HCMV replication. Gene expression and protein levels of host metabolic regulators and enzymes are altered within 48 hpi. ChREBP is a transcription factor regulated by glucose signaling and is activated by 24 hpi in HCMV infection (77, 78). The mature, active form of SREBP1 and SREBP2 transcription factors that promote cholesterol and fatty acid synthesis are increased 24 and 48 hpi, respectively (22, 79, 80). SREBPs help promote the expression of fatty acid elongase, ELOVL7, which is increased in HCMV infected cells by 24 hpi (8, 9, 78). HCMV infection promotes the levels of ACC1, the rate-limiting enzyme in fatty acid synthesis, by 48 hpi (9, 22, 79). Similarly, fatty acid synthase (FAS) is elevated by 48 hpi (22). Several kinases involved in HCMV reprogramming of metabolism are increased by 24 hpi, including CaMKK1 and PERK (10, 80, 81). While most of the viral mechanisms responsible for altering these host factors have yet to be elucidated, the kinetics of these alterations are consistent with an increase in de novo PC synthesis starting at 24 hpi (Fig. 2) and a shift in the PC lipidome between 24 and 48 hpi (Fig. 5). During this time new viral genomes are synthesized, which subsequently triggers the expression of HCMV late genes. Moreover, we found that PC changes induced by HCMV infection occur independent of late gene expression (Fig. 6).

After observing increases in the levels of PC lipids in several cell types under differing growth conditions following HCMV infection, we used metabolic labeling to determine how infection may alter the activity in PC synthesis pathways. Stable isotopic labeling using ^13^C, ^2^H or ^15^N tracers is a common approach for measuring intracellular metabolism. Most tracer studies performed to understand metabolic changes caused by virus infection have investigated metabolites or fatty acids (6–9, 66, 82). However, our study used metabolite and lipid tracers to analyze lipid labeling. We used labeling to determine which pathways are active in the fibroblast cells used for infection, which pathways are altered by infection, and when the pathways are altered. We found that HCMV infection increased the activity in the de novo synthesis pathway using ^13^C-choline (Fig. 2A). Moreover, ^13^C-choline labeling allowed us to determine when infection increased the de novo pathway. We found a marked increase in the de novo pathway starting at 24 hpi and persisting through at least 72 hpi (Fig. 2B-H). Additionally, by using a labeled lysoPC lipid tracer we were able to determine that LPCAT pathway is active in the fibroblast cells used in this study and that HCMV infection had little to no impact on the conversion of lysoPC to PC lipids (Fig. 4E-F). Finally, metabolic labeling allowed us to determine that the PEMT pathway that converts PE to PC was not active in HFFs and that HCMV infection does not promote its activation (Fig. 3). Overall, our observations using several metabolic tracers support the conclusion that HCMV infection increases the concentration of PC lipids by promoting the activity in the de novo PC synthesis pathway.

Our LC-MS/MS studies demonstrate the temporal changes to PC levels in HCMV-infection begin between 24 and 48 hpi and continue through late stages of infection (Figs. 2 and 5). Further, our data indicate that IE or E proteins induce metabolic reprogramming to feed and promote the de novo synthesis pathway of PC lipids. Overall, our study reveals that HCMV reprogramming of metabolism that supports a shift in the lipidome occurs independent of viral genome synthesis and without the need for late viral proteins. Knowing when metabolic changes are induced will allow us to better define how these changes occur and possible mechanisms to suppress the metabolic reprogramming that is essential for virus replication.

## MATERIALS AND METHODS

### Cells, viruses, and experimental design

HFF cells were used throughout the study. Where indicated, the study included additional fibroblasts (MRC-5 and MRC-9), ARPE-19 epithelial cells, and HUVEC endothelial cells. Fibroblasts were grown in DMEM, epithelial cells were grown in 1:1 DMEM:F12 medium, and endothelial cells were grown in Endothelial Cell Growth Medium (Lonza). Lipidomic labeling and replication experiments were performed in HFF-hTERT cells that were first grown to and held at 100% confluence for 3-days in Dulbecco’s modified Eagle’s medium (DMEM) with 10% fetal bovine serum (FBS), 10mM HEPES and penicillin/streptomycin (P/S). One day prior to HCMV infection, FBS was withdrawn from the medium. All HCMV infections were performed using TB40/E-GFP (9, 10). Cells were infected using a multiplicity of infection (MOI) of 3 infectious units per cell in serum-free (SF) DMEM, unless otherwise indicated. Mock-infected, uninfected cells received inoculum lacking virus particles and were otherwise treated the same in parallel with HCMV-infected cells. All infections were performed for 1h, after which the cells were washed with phosphate buffered saline (PBS). Cells were then maintained in their respective growth medias, unless otherwise stated. At 48 hpi, the growth medium was replaced to refresh nutrient levels, except for the PAA treatment conditions where cells were treated every 24h.

Experimental virus stocks were generated from a BAC-derived stock and propagated in fibroblasts at a low MOI. Extracellular virus from the supernatant was collected and concentrated to a pellet through a layer of sorbitol (20% sorbitol, 50mM Tris pH 7.2, 1 mM MgCl_2_) using ultracentrifugation at 20,000 rpm for 80 minutes. Virus was resuspended in SF DMEM with HEPES and P/S. Virus stocks and experimental samples were titered using a tissue culture infectious dose 50 (TCID_50_) assay. A starting dilution of 1:10 was used. After 2 weeks, infectious colonies were counted by the presence of two or more adjacent GFP^+^ cells.

### LC-MS/MS Lipidomics

Lipid abundances were measured using high-performance liquid chromatography high-resolution tandem mass spectrometry (LC-MS/MS). Briefly, cells were washed with PBS and treated with cold 50% methanol. Lipids were extracted twice using chloroform and dried under nitrogen (9, 10). Data were collected using either a Thermo Fisher Scientific Q Exactive Plus (QE+) or Thermo Fisher Scientific Exploris 240 mass spectrometer. For experiments performed on the QE+, lipids were resuspended in 100 µl of a 1:1:1 solution of methanol-chloroform-isopropanol per 200,000 cells. For experiments performed using the Exploris 240, lipids were resuspended in 200 µl 1:1:1 solution per 200,000 cells. For each sample, a total of three wells were used for analysis. Two wells were used for lipid extraction (i.e., technical duplicates), and one well was used to determine the total number of cells for normalization. Samples were normalized according to the number of live cells at the time of lipid extraction. Additional wells with no cells were used as a control to determine any contaminants from the lipid extraction and LC-MS/MS. Following resuspension, lipids were stored at 4°C or 7°C in an autosampler. Lipids were separated by reverse-phase chromatography using a Kinetex 2.6-mm C18 column (Phenomenex; 00F-4462-AN). LC was performed at 60°C using a Vanquish ultrahigh-performance LC (UHPLC) system (Thermo Scientific) and two solvents: solvent A (40:60 water-methanol, plus 10mM ammonium formate and 0.1% formic acid) and solvent B (10:90 methanol-isopropanol, plus 10mM ammonium formate and 0.1% formic acid). UHPLC was performed at a 0.25-mL/min flow rate, starting at 25% solvent B and ending at 100% solvent B as described previously (9, 10). After each sample, the column was washed and equilibrated. The total run time was 30 min per sample. Blank samples were run before, after, and interspersed with samples. Lipids were ionized using a heated electrospray ionization (HESI) source and nitrogen gas as described previously (9, 10). Data was collected using full scan MS/data-dependent MS2 (dd-MS2) TopN mode. QE+ MS1 abundance data were collected at a resolution of 70,000 over a mass range of 200 to 2,000 m/z and MS2 spectra were collected using a transient time of 128 ms and a resolution setting of 35,000 with an AGC target of 1e5. Exploris 240 MS1 abundance data were collected at a resolution of 90,000 over a mass range of 200 to 2,000 m/z and MS2 spectra were collected using a transient time of 64 ms and a resolution setting of 30,000 with an AGC target of 1e6. Each sample was analyzed using negative and positive ion modes. The mass analyzers were calibrated weekly and systems calibrated monthly.

### Lipid abundance analysis

PC abundance data were analyzed using MAVEN and a reference library of empirically derived retention times to aid identification of PCs (9, 10, 83). Lipids were initially identified in MS1 in positive mode using the precursor molecule’s calculated *m/z* and the anticipated isotopic pattern of naturally occurring ^13^C carbon. MS1 identifications were done using ≤8 ppm accuracy. In addition to high accuracy MS1 and retention time, the identification of all reported PCs was confirmed using MS/MS (MS2). In positive mode, PCs were confirmed by identifying the phosphorylcholine head group (184.0733 *m/z*). Moreover, all reported PCs were further confirmed in negative mode using formate-adducts. In negative mode, retention time, high resolutions MS1 accuracy, and tail identification were used to confirm the identity of PC lipids. All reported fatty acid (FA) tails were identified by MS2 in negative mode. Quantification of all PCs was done using MS1 peak information from positive ions.

### LC-MS/MS Metabolic Labeling Measurements

Stable isotope LC-MS/MS metabolic labeling experiments were performed as described above with the following differences in instrument settings and method. QE+ MS1 stable isotope tracer data were collected at a resolution of 140,000 and 280,000 resolution over a mass range of 400 to 1,400 m/z. Exploris 240 MS1 abundance data were collected at a resolution of 180,000 over a mass range of 200 to 2,000 m/z.

### Metabolic tracing

For stable isotope metabolic tracer studies, unlabeled PCs were identified using MAVEN and the predicted *m/z* of the precursor molecule (83). Labeled forms were exported to a csv. file from MAVEN based on the predicted *m/z* of the unlabeled precursor ion and mass shift produced by the isotopic tracer. MATLAB or RStudio IsoCorrectoR package were used to correct for the abundance of naturally occurring isotopes in the metabolic tracing experiments (6, 84). For *de novo* PC synthesis pathway analysis, the percentage of labeled PC was determined using 2-labeled ^13^C-choline which produces a MS mass shift of Δ2.006 *m/z* relative to unlabeled ^12^C-choline. Therefore, only 2-labeled PC forms possessing a 2-labeled choline head group from exogenously supplied ^13^C-choline were included. The amount of ^13^C-labeled PC was calculated for each species as the fraction of 2-labeled PC relative to the total measured PC level. For analysis of the PEMT pathway, ^13^C-methionine labeling was used. Since the PEMT pathway involves the sequential addition of three methyl groups to form a PC from a PE, the resulting PC will have up to three ^13^C carbon atoms depending on the labeling rate of S-adenosylmethionine (SAM) from methionine. Therefore, PCs containing 1-3 ^13^C atoms were included in the PEMT pathway analysis. The percent labeling was calculated as a fraction of the sum of 1-, 2-, and 3-labeled PC relative to the total PC level. For analysis of LPCAT activity, d5-LPC(17:0) containing 5 deuterium atoms in the glycerol backbone which produces a MS mass shift of Δ5.031385 *m/z* relative to unlabeled LPC(17:0) was used. Therefore, only 5-labeled PC forms were included in the LPCAT activity analysis. Candidate PCs were considered newly synthesized by LPCAT activity if they produced the anticipated Δ5.0313 mass shift and contained a mass spectral peak of 269.249 *m/z* corresponding to the C17:0 fatty acid of exogenously supplied d5-LPC(17:0). When present, the complementary PC tail was confirmed by FA tail analysis. The data was normalized following a normality test (RStudio) due to large differences in EIC values between label-fed vs EtOH control conditions, which was expected since d5-LPC(17:0) does not exist in untreated cells. An outlier test was used to determine that the low levels of background labeled PC(33:1) and PC(35:1) observed in 5 out of 24 EtOH samples were indeed outside of the normal distribution of the data. However, these data were included in the statistical analysis as their overall impact was relatively minor when weighed against the remaining samples, which exhibited no signal. A student t-test was used to determine statistical significance with a statistical threshold of *P<0.05*.

Metabolic tracer studies were performed using stable isotope labeling reagents purchased from Cambridge Isotope Laboratories Inc. and Avanti Polar lipids. ^13^C-choline (1,2,-^13^C2, Cambridge #CLM-548-0.1) was used to measure *de novo* PC pathway activity.^13^C-methionine (METHYL-^13^C, Cambridge #CLM-206-1) was used to measure PEMT pathway activity. LPC(17:0-d5) (AVANTI #855679) was used to measure LPCAT activity. Product quality reported >99.5% compound purity.

Growth medium containing either ^13^C-choline (4mg/L) or ^13^C-methionine (30mg/L) was generated following a published Gibco formulation (85). The solution was pH adjusted to 7.4 and filter sterilized. P/S and 10mM HEPES were added. All labeling experiments were performed in serum-free conditions as described above. The labeling media was stored at 4° C until used.

d5-LPC(17:0) in 1:1 dichloromethane-methanol was stored at −20° C until use. The day before use, the solvent was gently evaporated under nitrogen gas and was reconstituted using 100% EtOH; this was repeated for a total of 4 resuspensions. An aliquot of reagent from the same EtOH stock was used as the mock-treated control. The day of each experiment d5-LPC(17:0) was diluted into SF growth medium. Lipid-free BSA carrier protein (Sigma) was used at a final concentration of 1.7 µM to conjugate d5-LPC(17:0) prior to each experiment.

### PEMT cloning and overexpression

The human PEMT gene (NCBI CCDS ID #11187.1) was synthesized by GENEWIZ with a silent mutation at amino acid 27. The sequence was verified by next-gen sequencing. The 5’ sequence was engineered to contain an XbaI restriction site and the 3’ sequence was engineered to contain a ClaI site. XbaI-HF and ClaI restriction enzymes were used to cut the PEMT gene from the GENEWIZ vector. The gene was inserted into pLV-TRE-blast expression vector (VectorBuilder). HFF-hTERTs containing rtTA were transduced with pLV-TRE-PEMT lentivirus particles or pLV-TRE-GFP lentivirus particles as a control. Following blasticidine selection, PEMT or GFP overexpression was induced with doxycycline at the indicated concentration and validated by western blot.

### Protein analysis

Proteins were examined by western blot using SDS-PAGE. Proteins were resolved using Mini-Protean TGX 4-20% or anyKD gels (Bio-Rad) and transferred to an Odyssey nitrocellulose membrane (LI-COR). For PEMT detection, membranes were blocked using 1% BSA in phosphate-buffered saline with 0.05% Tween 20 (PBS-T) and incubated with a rabbit antibody against PEMT (Invitrogen PA5-42383) in 1% BSA PBS-T solution at 1:500 dilution. For all other blots, membranes were blocked using 1% milk in phosphate-buffered saline with 0.05% Tween 20 (PBS-T) and incubated with primary antibody in the presences of 1% milk-PBS-T solution. The following antibodies were used: rabbit polyclonal LPCAT1 (1:500; Cell Signaling E4V4B, #57411), mouse monoclonal IE1 (1:100; clone 1B12), mouse anti-UL44 (1:2,500; virusys), mouse anti-pp28 (1:100; clone 10B4-29), mouse anti-pp71 (1:100; clone 2H10-0), mouse anti-gB (1:50; clone 27-156), rabbit anti-tubulin (1:1,000; proteintech), and mouse monoclonal anti-α-tubulin (1:2,000; Sigma-Aldrich; #T6199). All HCMV antibodies except for anti-UL44 and anti-gB were gifts from Dr. Thomas Shenk (Princeton University). The anti-gB was provided by Dr. William Britt (University of Alabama at Birmingham) (PMID: 8614980). Membranes probed with mouse monoclonal antibodies were incubated for 1 h at room temperature (RT), while rabbit antibodies were incubated for 2 h at RT. Visualization and quantification of western blots were performed using a LI-COR Odyssey CLx imaging system.

### DNA quantification

Quantitative PCR (qPCR) was used to determine the HCMV genome copy number relative to cellular DNA. Cellular DNA was isolated using a Zymo Quick DNA mini-Prep kit (ThermoFisher Scientific). Viral genomes were quantitated between 4-96 hpi. Viral DNA levels were measured using primers specific to the HCMV gene UL123 (5’-GCCTTCCCTAAGACCACCAAT-3’ and 5’-ATTTTCTGGGCATAAGCCATA ATC-3’). Host genomes were measured using primers specific to actin (5’-TCCTCCTGAGCGCAA GTACTC-3’ and 5’-CGGACTCGTCATACTCCTGCTT-3’). A bacteria artificial chromosome (BAC) containing the viral genome sequence of HCMV FIX strain engineered to express cellular actin was used to develop a standard curve and determine the absolute quantities of viral DNA (86). qPCR was performed on a QuantStudio 3 Real Time PCR system using PowerUp SYBR Green Master Mix (ThermoFisher Scientific).

### Statistics

Figures and graphs were generated using GraphPad Prism and RStudio. Statistical significance was determined for Figure 1B-C using a t-test with a significance cutoff of *P <0.05*. Figures 2 and 6 use a Two-way ANOVA and Tukey’s test for post-hoc analysis.

## ACKNOWLEDGMENTS

We want to thank Dr. Roslyn Dermody and the other members of the Purdy and Goodrum laboratories for assisting with this study. This project was supported by the National Institute of Health (NIH) National Institute of Allergy and Infectious Disease (NIAID) R01AI162671 (J.G.P.), R01AI155539 (J.G.P.), F32AI178919 (R.L.M.), and National Institute of Aging (NIA) T32AG058503 award (I.K.). The content is solely the responsibility of the authors and does not necessarily represent the official views of the NIH. Additional funding was provided by the BIO5 Institute Postdoctoral Fellowship program awarded to R.L.M. and by the Personalized Defenses Against Disease strategic initiative from the University of Arizona.

## AUTHOR CONTRIBUTIONS

I.K., R.L.M., Y.X., and J.G.P. conceptualized and designed experiments; I.K., R.L.M., Y.X., M.A.M., N.S., S.L., and J.G.P. performed experiments. I.K., R.L.M, and M.P.M. analyzed the data. I.K., R.L.M., M.P.M., N.S., S.L., and J.G.P. interpreted the results; I.K. and J.G.P. drafted the manuscript; All authors reviewed and edited the manuscript.

